# The contrasting flowering-time among coffee genotypes is associated with ectopic and differential expressions of genes related to environment, floral development, and hormonal regulation

**DOI:** 10.1101/2024.08.02.605191

**Authors:** Marlon Enrique López, Raphael Ricon de Oliveira, Lillian Magalhães Azevedo, Iasminy Silva Santos, Thales Henrique Cherubino Ribeiro, Dapeng Zhang, Antonio Chalfun-Junior

## Abstract

The molecular pathways underlying floral activation and development are well described in model species, but exhibit significant diversity in plants that is poorly understood in crops with complex cycles, such as *Coffea arabica* L. The reproductive development of coffee plants is biannual, and the flowering time is crucial for the coffee productivity and cup quality. In this study, we explored the plasticity of floral development and flowering-time of contrasting coffee genotypes to understand the associated metabolic and regulatory transcriptional profiles. Firstly, we compared the reproductive development of three coffee genotypes, confirming that *Acauã* is late flowering, *Oeiras* is early flowering, and the natural mutant *Semperflorens* (*Sf*) exhibits continuous flowering throughout the year. Analysis of sugar and ethylene content revealed quantitative differences between genotypes in both leaves and floral buds. To associate these phenotypic differences with the regulatory developmental pathways, we performed RNA-seq analysis comparing the shoot apical meristems, floral buds and leaves of different genotypes. Our analysis identified 12.478 differentially expressed genes, which showed enriched terms mainly related to hormonal regulation, external stimulus and floral development. Notably, some major players of reproductive development, as homologs of *FLOWERING LOCUS T* and MADS-box genes, showed contrasting expression patterns, generally being ectopically upregulated in the *Sf* mutant. These findings were associated with the phenotypic differences among coffee genotypes. In conclusion, the present study improves the understanding of the divergence of floral development in coffee, providing valuable insights for directing breeding programs and future studies aiming at controlling floral development and enhancing crop production.

## 1. INTRODUCTION

The flowering process is intricately regulated by multiple external and endogenous signals to timing the transition of meristems from vegetative to reproductive stages (Coneva et al., 2012). Numerous pathways are described regulating floral development, such as the well described photoperiod, gibberellins, vernalization, aging and the autonomous pathways (Ausin et al., 2005; Blümel et al., 2015). Additionally, sugar status, stress and other plant hormones play roles in this process (Izawa, 2021). All these pathways are interconnected and divergent between plants, reflecting the different reproductive strategies and a complex relationship (Blümel et al., 2015; Krizek & Fletcher, 2005). Understanding these pathways and its connection with the flowering process is particularly challenging in Arabica coffee (*Coffea arabica* L.), a woody species that presents a biennial cycle regulated by external and endogenous stimulus and an allotetraploid genome (reviewed by López et al., 2021).

The flowering pattern of coffee is classified as gregarious, meaning anthesis occur simultaneously within a certain geographical localization, but with different numbers of flowering events depending on the drought and rainfall regime (Cannell, 1985; Crisosto et al., 1992; Ronchi et al., 2015). This asynchronous flowering leads to varied starting points of fruit development, resulting in uneven fruit production at the harvesting time, which impairs the yield and coffee cup quality (Cannell, 1985; Lima et al., 2021). Studies have been carried out to synchronize coffee flowering events, aiming to improve homogeneity during fruit ripening (Lima et al., 2021; Miranda et al., 2020). Other factors, such as photoperiod, temperature, shade conditions, plant nutritional status, and phytohormones can also affect the floral transition and the coffee reproductive cycle (López et al., 2021; Ramírez Builes et al., 2013).

Recently, it was observed that the dry period following rehydration in coffee trees is associated with gene regulation of ethylene biosynthesis and signaling pathway. This promotes anthesis by increasing ethylene level in the shoot and, by increasing ethylene sensitivity through receptors such as *CaETR4-like* (Lima et al., 2021; Santos et al., 2022). Similarly, ethylene biosynthesis pathway induction positively regulated natural flowering in pineapple (Trusov & Botella, 2006). However, in rice, analysis of the ethylene receptor gene *ETR2* revealed that ethylene signals delay floral transition (Wang et al., 2013; Wuriyanghan et al., 2009)). Endogenous regulation of other hormones levels is also associated with flowering regulation, such as brassinosteroids that promote a floral transition in *A. thaliana* (Z. Li & He, 2020), and abscisic acid (ABA) that affects a floral transition under water deficit conditions in rice (Du et al., 2018). In coffee trees, ABA increases during the dry period and is associated with floral bud latency of the G4 state, whereas plant rehydration promotes decreasing of ABA (López et al., 2021) and increasing of gibberellin (GA) levels (Browning, 1973). Growth regulators can also promote flowering, such as exogenous cytokinin (D’Aloia et al., 2011), gibberellin (Schuch et al., 1990), and the ethylene inhibitor 1-methylcyclopropane (1-MCP) (Lima et al., 2021).

Another important factor described as a flowering regulator is carbohydrates. Sucrose level increased during the transition of the vegetative to reproductive phase, acting as a signal to control development (Horacio & Martinez-Noel, 2013). The higher energy demand in the reproductive system, driven by processes such as mitosis and the formation of flowers, seeds and fruit organs, underscores the roles of sucrose in the flowering transition (Bernier et al., 1993).

Despite evidence supporting the participation of sugar in signaling the flowering process, the mechanism of how it participates and transmits signals remains largely unknown. Previous studies reported that sucrose has the main function in the leaf phloem for the generation of florigens such as the FLOWERING LOCUS T (FT), whereas trehalose-6-phosphate acts in the shoot apical meristem to promote the flowering signal of these florigens (Cho et al., 2018). Furthermore, sugar transporters are involved in the long-distance movement of sucrose from source to sink, a critical process for phloem loading (Smeekens et al., 2010). Similar to the hormonal crosstalk associated with flowering regulation, the relationship of sugars with flowering is not universally conserved, with varied responses observed in different species. Exogenous application of sucrose has shown floral induction followed by a high expression of *FT* homologs, in Chrysanthemum (*Chrysanthemum indicum*) confirming the signaling functions of sucrose in this species (Sun et al., 2017). Conversely, the application of 6% of glucose delays flowering in *Arabidopsis* associated with Long Day (LD) photoperiod (Zhou et al., 1998).

While much research has focused on elucidating the role of sugars in grain development and its effect on the coffee cup quality (Privat et al., 2008; Rogers et al., 1999; Torrez et al., 2023), limited information exists on how sugar metabolism influences the flowering process in coffee trees. Despite the conserved pathways activating the floral development, i.e. the florigen *CaFT1* in *C. arabica* (Cardon et al., 2022), the relationship with the sugar status, hormones and environmental cues remains unclear.

In this study, we hypothesized that differences in the floral development among coffee genotypes might be associated with the sugar metabolism and differences in transcriptional profiles at key stages of development. To explore this hypothesis, we comparatively analyzed three genotypes (*Oeiras*, *Acauã* and the mutant *Semperflorens*), respectively described as early-, late-and continuous-flowering (Antunes, 1960; Carvalho & Krug, 1952). Firstly, we evaluated the phenotypic differences along the reproductive cycle of the three genotypes, then we performed sugar and ethylene analysis and transcriptional analysis through RNA Sequencing (RNA-seq) of the shoot apical meristem (SAM), floral buds (FBs) in the beginning of development -G2 stage (Morais et al., 2008), and leaves collected in March and August to represent the contrasting environmental conditions of Brazil (cold/dry vs warmer/raining). The contrasting reproductive development of coffee genotypes were associated with metabolic and transcriptional patterns, mainly related to environment-, floral-and hormonal-pathways. Thus, this study contributes to understanding the divergence of floral development in crops with complex developmental cycles, providing insights for directing breeding programs towards the control of coffee reproductive development and enhancing coffee production.

## 2. MATERIAL AND METHODS

### 2.1. Plant Material

Three *Coffea arabica* genotypes were chosen based on their contrasting reproductive development patterns: cvs. *Oeiras* and *Acauã*, representing early-and late-flowering respectively, and the natural coffee mutant *Semperflorens* (*Sf*) that exhibits a continuous-flowering phenotype (Antunes, 1960; Carvalho & Krug, 1952; Carvalho et al., 2008). In details, *Acauã* (IBC – PR 82010) comes from the cross between ’Mundo Novo IAC 388-17’ and ’Sarchimor IAC 1668’, whereas *Oeiras* is derived from the hybrid CIFC HW 26/5 after the crossing between ’Caturra Vermelho’ (CIFC 19/1) and ’Híbrido de Timor’ (CIFC 832/1), resulting in the *Catimor* genotype (Carvalho et al. 2008).

*Sf* is a natural mutant that flourishes continuously throughout the year due to a mutation occurring in a “Bourbon” genetic background. However, the mutated regulatory mechanism responsible for its phenotype has not been elucidated (Antunes, 1960; Carvalho & Krug, 1952; Carvalho et al. 2008). The experiment was conducted in 2020-2021 harvest season using five-year-old plants established at the experimental field of National Institute of Science and Technology (INCT-Café) located at the Federal University of Lavras (UFLA; Lavras, Minas Gerais/Brazil, 21°23′S, 44°97′W). In this field, ten plants of each coffee genotype were grouped in different blocks, and such blocks were spatially distributed randomly, with the plants being cultivated following the standard recommendations for nutritional and pest control (Vieira, 2008).

The plant material used for analyses consisted of four biological replicates, each with three plants, totaling twelve plants per genotype and time point. Plant sampling consisted of leaves (fully expanded) or FBs (at different developmental stages) pooled and collected monthly from March to September 2020 and shoot apical meristems (SAM) collected in January 2021, between 4:00 to 5:00 p.m. Samples for carbohydrates and RNA-seq analyses were immediately immersed in liquid nitrogen and stored at -80 ◦C until use. Materials for ethylene analysis were incubated directly in vacuum glass tubes. A detailed summary of the tissue sampling and collection dates are available in Fig. S1.

### 2.2. Analyses of floral bud development and flowering time (anthesis)

To compare the floral development patterns and flowering time across coffee genotypes, the number and developmental stage of FBs in the nodes of plagiotropic branches were evaluated monthly until anthesis (opening of flowers). In details, for each coffee genotype it was used three plants per biological replicate and marked six nodes in four plagiotropic branches per plant (3 plants x 4 plagiotropic branches x 6 nodes = 72 nodes per biological replicate), totaling 288 nodes analyzed for each coffee genotype spanning four biological replicates. The number of FBs were quantified, and their developmental stages determined according to Morais et al. (2008) monthly from April till June 2020 (Fig. 1A). The proportion of opened flowers and number of anthesis events were also quantified using three plants per biological replicate but selecting three nodes of four plagiotropic branches per plant (3 plants x 4 plagiotropic branches x 3 nodes = 36 nodes per biological replicate). This analysis covered three biological replicates, totaling 108 nodes per genotype and extended from March to September 2020 (Fig. 1B), with the final period coinciding with the rainy season in Brazil.

**Fig. 1.**
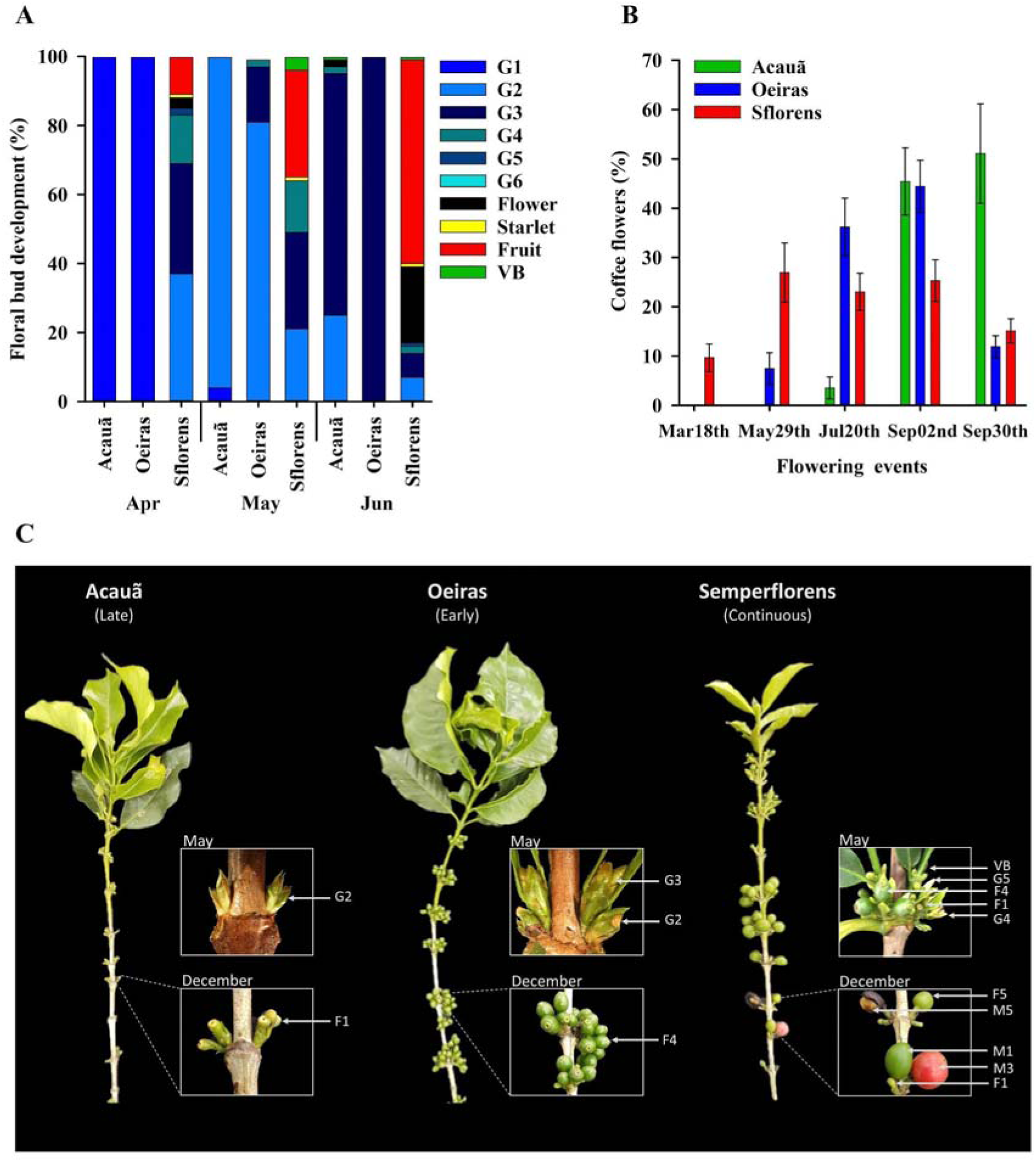
The floral development patterns and flowering-time of three contrasting coffee genotypes. The patterns of reproductive development were determined in the *Acauã* and *Oeiras* genotypes described as late-and early-flowering, respectively (Carvalho et al. 2008), and in the natural mutant *Semperflorens* (*Sf*) that flourishes continuously throughout the year (Antunes, 1960; Carvalho & Krug, 1952; Carvalho et al. 2008). **A-** Figure shows the proportion of FBs and their developmental stages in April (Apr), May and June (Jun). **B-** Figure shows the proportion of opened flowers at different months (flowering events) comparing coffee genotypes. **C-** Details of the differential reproductive development found in May and December between genotypes. Legends: G1 to G6 refers to the progressive stages of floral bud development, F1 to F5 stages of fruits development and M1 to M5 stages of fruit, all of them described in Morais et al. (2008); VB -vegetative buds.

### 2.3. Carbohydrates and ethylene content

Extraction of carbohydrate followed the methods adapted by Silva et al. (2014) but using 200 mg of fresh frozen tissue (completely expanded leaves; Fig. S1) previously powdered with liquid nitrogen and homogenized in a 5 mL potassium buffer (100 mM, pH 7.0). Starch, sucrose, and total soluble sugars were quantified according to Dische (1962), and reducing sugar levels according to Miller, (1959). Ethylene quantifications were made using leaves and floral bud tissues collected at different time points (see Fig. S1 for details), following the method described in López et al. (2022), with the production rate expressed as ppm. g^-1^ FW. h^−1^ (Fresh weight per hour).

### 2.4. RNA extractions and RNA sequencing (RNA-seq) libraries preparation

Total RNA from leaves, FB, and SAM were extracted according to de Oliveira et al., (2015). RNA integrity was checked in agarose gel (1%) whereas the purity and composition were accessed by spectroscopy (NanoVue GE Healthcare). All the samples presented OD^260/280^ and OD^260/230^ *>* 1.8 and <2.1. Next, 7.5 μg of RNA were treated with DNase I (Turbo DNA-free Kit, Ambion) following the manufacturer’s instruction to eliminate DNA contamination, and the RNA integrity numbers (RIN) checked again by Agilent 2100 Bioanalyzer (Agilent Technologies), in which all the samples presented standard values higher than 6.0.

A total of 2,765,320,286 of non-strand-specific paired-end reads (2 x 150 bp) were sequenced using an MGI-tech DNBSEQ-G400 sequencer. The tissues used were obtained from leaves (March or August/2020), FBs and SAM tissues of the three coffee genotypes in four biological replicates, totalyzing 48 RNA-seq libraries (details in Fig. S1). The sequencing libraries and quality control were done by BGI Genomics Co., Ltd. The method for filtering coding RNAs from other classes of RNAs was the polyA selection using oligo dT. Further, parameters such as overall sequence quality, sequence length distribution, duplication levels, overrepresented sequences, and adapter content were accessed using FastQC version 0.11.8 (Wingett & Andrews, 2018). Raw data for the RNA-seq runs can be obtained through BioProject ID PRJNA1091177 and the respective SRA accessions are available in Table S1.

### 2.5. RNA-seq and Differential Expression (DE) analyzes

RNA-seq libraries were aligned to the *C. arabica* genome (available at https://www.ncbi.nlm.nih.gov/genome/?term=txid13443) using the annotation Cara_1.0 with STAR version 2.7.8a (Dobin et al., 2013). The resulting alignment files were sorted and inspected with Picard toolkit (version 2.25.4, 2019) to remove all duplicated fragments that otherwise would inflate counts and bias subsequent analysis (Peng et al., 2015). Fragments uniquely mapped to exons were quantified with the HTSeq-count script (Anders et al., 2015) and the number of sequenced fragments per sample, uniquely mapped, multi mapped, or unmapped to the *Coffea arabica* genome are available in Fig. S2.

After quantification, fragment counts were summarized with in-house R scripts and normalized using edgeR (GLM mode) from the Bioconductor package (Huber et al., 2015; Robinson et al., 2010). This procedure allowed us to infer the relationship between different samples and gene expressions in relevant library contrasts, throughout a Multidimensional Scaling (MDS) plot (Fig. S3-A), a heatmap representation of the Spearman correlation (Fig. S3-B) and a hierarchical cluster analysis on the set of dissimilarities of Differentially Expressed (DE) genes across the evaluated conditions representation of the relationships between all differentially expressed genes and conditions (Fig. S3-C). More details of this analysis are available in the legend of figures. Genes were considered DE when their false discovery rate -FDR (Benjamini & Hochberg, 1995) - were below 0.05 and had a minimum expression fold change of two. In other words, a given gene was called DE if the difference in expression between two conditions is at least two fold increase the expression in the condition with the lower value and passed an FDR threshold.

### 2.6. Statistical analysis

For sugars and ethylene content data analyses were performed using the InfoStat software (Di Rienzo et al. 2019). The statistical difference was determined by one-way ANOVA, followed by the Tukey test. Results were expressed as the mean ±Standard Deviation (SD). Details are available in Table S2.

## 3. RESULTS

### 3.1. Coffee genotypes exhibit different patterns of floral development and flowering time

To explore the underlying molecular pathways regulating floral development, we initiated by characterizing the contrasting floral development in the selected coffee genotypes: cvs. *Acauã* and *Oeiras* described as late-and early-flowering, respectively (Carvalho et al. 2008), and *Sf*, a natural mutant that flourishes continuously throughout the year (Antunes, 1960; Carvalho & Krug, 1952; Carvalho et al. 2008). The number and developmental stage of floral buds (FBs) were estimated and compared between genotypes in different months until anthesis and fruit formation (Fig. 1). The G1 stage corresponds to the first observable stage of buds development, when it protrudes outside the interpetiolar stipules becoming visible and develops until G6, the last before anthesis (de Oliveira et al., 2014; Morais et al., 2008).

In April, both *Acauã* and *Oeiras* displayed all FBs at G1 whereas the *Sf* presented a mix of different stages (including fruits), which aligns with its typical phenotype of continuous flowering (Fig. 1A). By May, differences in the floral development progression became evident with *Acauã* presenting most of the FBs at G2 whereas *Oeiras* displayed a considerable number of FBs at G3 (Fig. 1A). This pattern persisted into June, where *Acauã* continued to have FBs at G2, and mostly at G3, while all FBs in *Oeiras* were at late G3 stage, representing an accelerated floral development process (Fig. 1A and B) consistent with its reported early flowering phenotype (Carvalho, 2008). In detail, both genotypes presented four anthesis events being two majors with a higher number of opened flowers (Fig. 1B). Whereas *Sf* showed five anthesis events distributed throughout the year in a different pattern in comparison with the other genotypes and the unique flowering event reported in March (Fig. 1B).

The differential development patterns of FBs described above (Figs. 1A and B) are summarized in Fig. 1C, illustrating the typical reproductive phenotype of each genotype. In May, the late-flowering cv. *Acauã* presents FBs in G2, while the early-flowering *Oeiras* in G2 and G3. On the other hand, *Sf* has FBs in more advanced stages (G4 and G5), accompanied by fruits (F1 and F4). After anthesis in December, *Acauã* presented F1 fruits, *Oeiras* already had F4 fruits, while *Sf* presented fruits at various stages (Fig. 1C). Thus, these results agree with the described characteristic of each genotype, confirming the contrasting development and flowering-time between them and suggesting differences in the regulatory molecular pathways, which we further explored.

### 3.2. Carbohydrates and ethylene quantifications reveal contrasts between coffee genotypes supporting differences in the reproductive development

Since floral development and flowering time (anthesis) are both related to sugar metabolism (Bolouri Moghaddam & Van den Ende, 2013; Cho et al., 2018; Ohto et al., 2001); and ethylene biosynthesis (Achard et al., 2007; López et al., 2022), the profile of these metabolics were evaluated in the leaves and floral buds of the three contrasting coffee genotypes at different periods of the year (Fig. 2). Analysis of starch content reveals a gradual increase from March to August in all genotypes, but with quantitative differences between them in some periods (Fig. 2A). Specifically, *Sf* always presented higher levels of starch compared to other genotypes, except in September, reaching the highest level in July, while *Acauã* and *Oeiras* showed higher levels in August. Regarding sucrose, the genotypes *Oeiras* and *Sf* presented similar accumulation patterns throughout the period, differing only in March, whereas *Acauã* showed higher levels than the other genotypes in June to August (Fig. 2B). The statistical analysis supporting these results is available in Table S2.

**Fig. 2.**
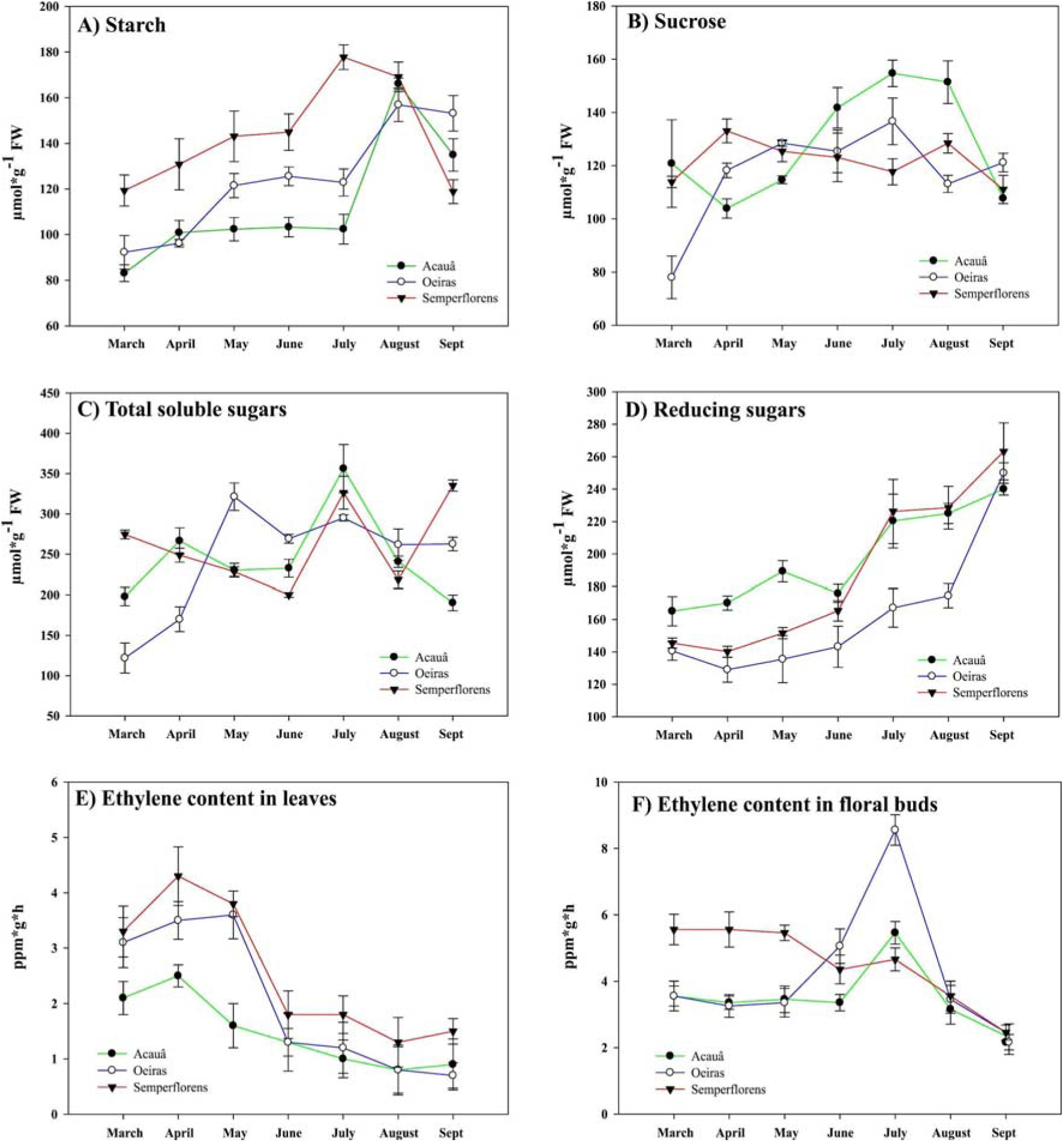
Carbohydrates and ethylene contents in tissues of coffee genotypes from March to September. Analyses were made in leaves and floral buds of the genotypes *Acauã* (late-flowering), *Oeiras* (early-flowering) and *Semperflorens* (continuous flowering) at different seasons in Lavras/MG (21°13’35.1"S 44°58’14.4"W, Brazil). This municipality is situated at approximately 919 meters above sea level with the dry/cold period extending from June to August while the rainy/warm period from September to May. Details of the local subtropical climate are available in (Dantas et al., 2007). The content of Starch (A), Sucrose (B), Total soluble sugars (C), Reducing sugars (D) were measured in leaves, whereas the ethylene production in leaves (E) and in floral buds (F). Legends: Mar - March, Apr - April, Jun - June, Jul - July, Aug - August, Sep - September.

Analysis of other carbohydrates such as total soluble sugars (TSS) showed a very similar pattern in *Acauã* and *Sf,* with the highest level for both in July, but they were different in March and September, being higher in *Sf* (Fig. 2C). This pattern was different from *Oeiras* that presented an TSS increase from March to April, becoming stable afterwards (Fig. 2C). The levels of reducing sugars (RS) presented a gradual increase from March to September for all genotypes, differing only quantitatively, with *Acauã* presenting higher levels in March to June, while *Oeiras* had lower levels in all timepoints (Fig. 2D). Details of statistical analysis are available in Table S2.

The quantification of ethylene in leaves revealed similar patterns for all genotypes, with the highest levels being reached in April and decreasing progressively until September (Fig. 2E), although *Acauã* showed lower levels in all timepoints. On the other hand, for floral buds the pattern of ethylene levels was only similar between *Acauã* and *Oeiras*, reaching the highest level in July, which contrasted with *Sf,* showing a pattern of higher levels in March to May decreasing progressively until September (Fig. 2F). In addition, comparing genotypes, *Oeiras* showed the highest level of ethylene in floral buds closely to anthesis (Fig. 1F), which is in agreement with the ethylene production and sensitivity associated at the period (Lima et al., 2021) and also its early-flowering phenotype (Fig. 1) (Carvalho et al. 2008).

Altogether, these results showed that coffee genotypes exhibit different profiles of primary metabolism throughout the plant developmental cycle (Fig. 2). These variations in chemical content coincide with observed differences in flowering developmental timing (Fig. 1). After showcasing the phenotypic and metabolic variations among coffee genotypes, we subsequently explored the molecular pathways that underlying these developmental differences.

### 3.3. Gene expressions differed mainly in function of organs rather than genotypes, but not for the *Sf* mutant which showed particular characteristics

Out of all sequenced reads, approximately 2 billion were uniquely mapped to exons in the genome - or 74.8% of the total (the mean values of fragments mapped or un-mapped per library are available in Fig. S2). After removing duplicated fragments, a total of 876 million high-quality individual fragments, representing a mean value of 18.3 million fragments in each library were uniquely mapped to exons in the *C. arabica* genome (Effective Counts in Fig. S2). These quantification matrices were then used for subsequent Differential Expression (DE) analyses. Finally, a total of 12,478 genes, 15% of all annotated in the reference genome, were found to be DE in the analyzed conditions (Figure S3) and their expression profiles are available in the supplementary file Barplots.

The quantified fragments were normalized to infer the general relationships between RNA-seq libraries and their differentially expressed genes (Fig. S3). First, a Multidimensional Scaling (MDS) was used to access the Euclidean distance and compare the expressed genes in each pair of samples (Fig. S3-A). In other words, the similarity between RNA-seq libraries was determined, where closer distance indicated more similar expression of the genes in the samples (Huber et al., 2015; Robinson et al., 2010). As expected, samples of the same organ clustered together independently of the coffee genotype (Fig. S3-A), i.e., all leaves and shoot apical meristems (SAMs), reflecting the similarity of biological replicates. However, considering the floral buds (FBs) samples, the library’s clustering was observed for *Oeiras* (Oei) and *Acauã* (Aca) genotypes, but not for the *Sf* mutant, which grouped separately. Moreover, the FBs RNA-seq libraries of *Sf* (“Sem FBuds” in Fig. S3-A) were closer to the SAM cluster than the FBs of other genotypes (“Oei and Aca FBuds” in Fig. S3-A). This result supports the differential flowering phenotype of *Sf* and suggests that its continuous flowering could be related to the maintenance of less differentiated meristems (more similar to SAM) within FBs (Fig. S3).

In addition, two other analyses were made to calculate the correlation of normalized gene expressions and RNA-seq libraries using the Spearman method of Hierarchical Clustering Tree (Fig. S3-B and C). Both figures, one comparing RNA-seq libraries to each other (Fig. S3-B) and the other individual DE genes per library (Fig. S3-C), showed three clear distinctive clusters: 1) all SAM samples grouped together in addition to the FBs libraries of the *Sf* mutant, 2) FB libraries from cvs. *Oeiras* and *Acauã* and, 3) another composed of leaves samples from all genotypes. Interestingly, leaves libraries of cv. *Acauã* and *Oeiras* grouped according to the collection period (March or August), but for *Sf*, the libraries grouped together independently of the period. In other words, the gene expression in RNA-seq libraries of leaves were similar between *Acauã* and *Oeiras* genotypes considering one specific period, but the *Sf* libraries showed more similarity between leaves of August and March, not distinguishing by period (“SemLeafAug_20” in Fig. S3-C). These results were similar to the MDS analysis (Fig. S3-A) showing that gene expressions differed mainly in function of organs. However, the *Sf* mutant showed unique characteristics regarding gene expression: 1) FB libraries closer to SAM, which suggest FBs together meristem-like tissues, and 2) leaves of different time points (March and August) closely related, suggesting a transcriptional homeostasis during different periods and/or a reduced sensitivity to environmental cues, given that March represents the rainy season while August corresponds to a drought period in Brazil.

### 3.4. An overview of the transcriptional patterns indicated differentially expressed genes (DEGs) and similarities between coffee tissues and genotypes

To obtain an overview of the molecular pathways (de)activated during development of the contrasting coffee genotypes, we compared the differentially expressed genes (DEGs) between tissues, selecting eighteen relevant comparisons (Table S3, Figs. S4). It is important to note that the reference in each comparison is the second RNA-seq library (after the “_x_”) pointing out which DEG is down-or upregulated in relation to the other library. The total number of DEGs between FBs and SAM tissues is supposed to indicate genes related to the floral identity (SAM is the progenitor of floral meristems and a less differentiated tissue).

These DEGs are more abundant in the coffee genotypes *Oeiras* and *Acauã*, 8143 and 7454 respectively, compared to the *Sf* mutant with 3946 (Table S3, Fig. S4-A to C). This agrees with the previous result (Fig. S3-A), in which the SAM and FBs libraries of the *Sf* mutant grouped closely, perhaps explaining its capacity to generate FBs continuously throughout the year.

By comparing identical organs between genotypes, we show that SAM libraries are similar regarding the abundance of DEGs for *Acauã* vs *Oeiras* and *Oeiras* vs *Sf*, 1226 and 1097 respectively (Table S3 and Figs. S4-D and F). However, the abundance of DEGs for *Acauã* vs *Sf* is lower, with 479 being DE between the genotypes (Table S3 and Fig S4-E). This result was interpreted as the SAM being transcriptionally more similar between *Acauã* and *Sf* mutant than *Oeiras*, which could be related to their genetic background since *Acauã* and *Sf* are closer related (details in “Plant Material” section). For FBs libraries, the comparisons pointed out that *Acauã* and *Oeiras* are transcriptionally more similar, 937 DEGs (Table S3 and Fig. S4-G), since both genotypes presented a higher number of DEGs compared to *Sf*, 5647 and 6700 respectively (Table S3, Fig. S4-H and I). This confirmed that the floral transcriptional pathways of *Sf* are different in accordance with its atypical continuously flowering phenotype (Antunes, 1960; Carvalho & Krug, 1952; Carvalho et al. 2008).

The other nine comparisons were made similarly as described above but contrasting RNA-seq libraries of leaves collected in March and August (Table S3 and Fig. S5; details of material in Fig. S1 and “Plant Material” section). Within a given genotype but at distinct time points, *Oeiras* and *Acauã* exhibited 3866 and 3666 DEGs, respectively (Table S3, Fig. S5-A and B) whereas *Sf* demonstrated 1524 DEGs (Table S3 and Fig. S5-C). These results indicated the leaf transcriptional profile comparing March (the rainy and warm period) to August (the dry and cold period) is less different for the *Sf* mutant, suggesting homeostasis or a greater insensitivity to environmental factors than the other genotypes.

When comparing libraries from leaves collected in the same months but representing different cultivars, a total of 839 differentially expressed genes (DEGs) were identified in the *Acauã* vs *Oeiras* comparison and documented in Table S3 and Fig. S5-D. Similarly, the *Acauã* vs Sf comparison revealed 1961 DEGs (Table S3 and Fig. S5-E), while the *Oeiras* vs Sf comparison exhibited 2073 DEGs (Table S3 and Fig. S5-F) during the month of March. In the month of August, the *Acauã* vs *Oeiras* comparison demonstrated 2908 DEGs (Table S3 and Fig. S5-G), the *Acauã* vs Sf comparison displayed 3413 DEGs (Table S3 and Fig. S5-H), and the *Oeiras* vs *Sf* comparison showed 4023 DEGs (Table S3 and Fig. S5-I). This result suggests that during August, the cultivars tend to rely on more distinct transcriptional strategies to cope with the environment.

### 3.5. Floral development and leaf function are mainly regulated by ethylene-and auxin-related pathways

To elucidate the significance of differentially expressed genes (DEGs) across the eighteen comparisons (detailed in Table S3 and Figs. S3, S4 and S5), we conducted a comprehensive Gene Ontology (GO) enrichment analysis. This involved categorizing the groups of up-or down-regulated DEGs in terms of their molecular function, biological process, and cellular component (Suppl. files “GO”). These analyses resulted in complex relationships with many significant GO terms featuring at multiple interconnected levels such as reproductive development and response to external stimulus (Suppl. files “GO”). Then, based on the main described pathways related to flowering regulation (Izawa, 2021; Kim, 2020; Leijten et al., 2018), we selected eleven enriched and relevant GO terms categories to classify the DEGs of the eighteen contrasts: Circadian Clock (CC), Endogenous Stimulus (EnS), External Stimulus (ExS), Floral Transition (FT), Floral Development (FD), Hormonal Regulation (HR), Photoperiod (PP), Photosynthesis (PS), Sugar Metabolism (SM), Temperature (T) and Unknown (U; Figs. 3 and 4). In our analysis, we observed distinct sets of up-and down-regulated genes within each GO term for every comparison. However, certain terms, such as ExS, FD, and HR, stood out for their significantly higher levels of enrichment, as illustrated in Figures 3 and 4.

**Fig. 3.**
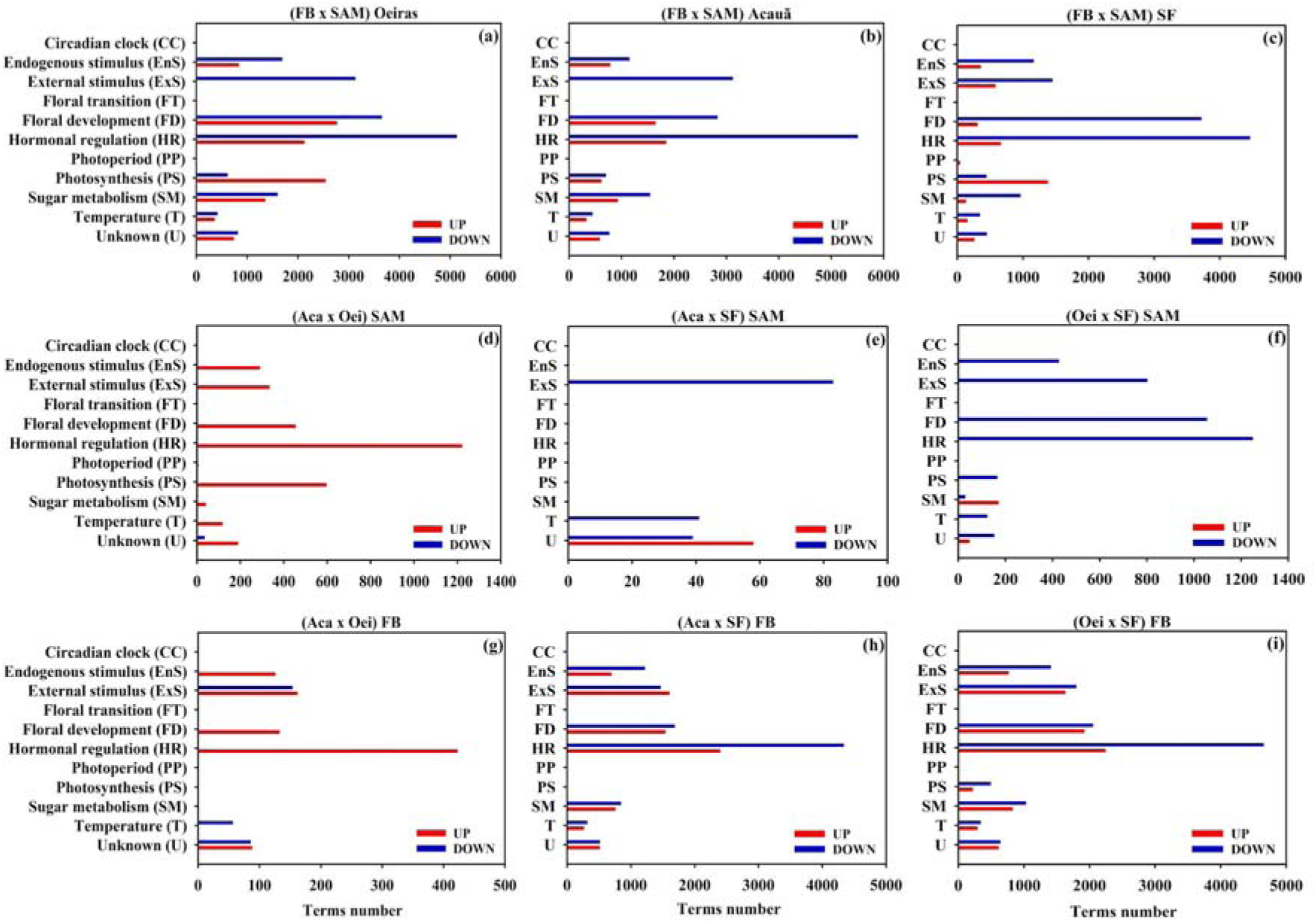
The number of selected Gene Ontology (GO) terms related to UP and DOWN-Regulated genes found in tissues of the coffee genotypes. The three upper figures show the contrast comparing different tissues, such as Floral bud (FB) and Shoot Apical Meristem (SAM), of the same genotype: *Oeiras* (Fig. a), *Acauã* (Fig. b) and *Semperflorens* mutant (Fig. c). The contrasts comparing the same tissue but different genotypes are shown in Figs. d to i. Legends: FB, Floral buds at the G2 stage; SAM, Shoot Apical Meristem; Aca, *Acauã*; Oei, *Oeiras*; SF, *Semperflorens* mutant; UP (in red), number of terms related to UP-regulated genes; DOWN (in blue), number of terms related to DOWN-regulated genes. Details of the collected tissues and periods are available in Fig. S1.

**Fig. 4.**
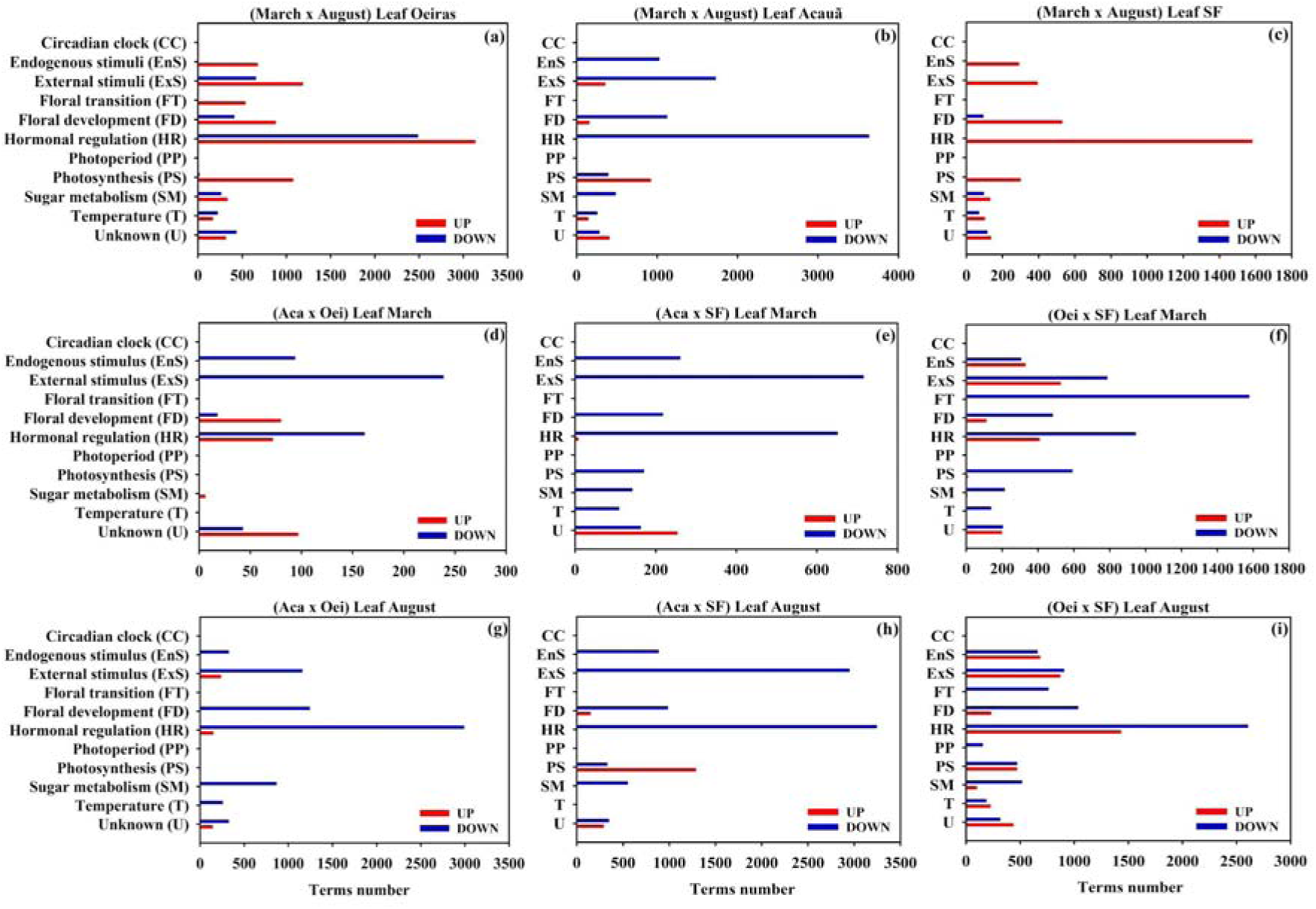
The number of selected Gene Ontology (GO) terms related to UP and DOWN-Regulated genes found in leaves collected in different periods of coffee genotypes. The three upper figures show the contrast comparing leaves collected in March or August of the same genotype: *Oeiras* (Fig. a), *Acauã* (Fig. b) and *Semperflorens* mutant (Fig. c). The contrasts comparing leaves collected at the same periods but different genotypes are shown in Figs. d to i. Legends: Aca, *Acauã*; Oei, *Oeiras*; SF, *Semperflorens* mutant; UP (in red), number of terms related to UP-regulated genes; DOWN (in blue), number of terms related to DOWN-regulated genes. Details of the collected tissues and periods are available in Fig. S1.

In detail, comparing FB to SAM libraries, all coffee genotypes presented similar patterns with a higher number of GO terms of DEGs (both UP and DOWN-regulated genes) related to ExS, FD and HR in relation to the SAM (Fig. 3A to C). Comparing SAM libraries between genotypes, *Oeiras* showed up-regulated genes compared to *Acauã* mainly related to FD, HR and PS (Fig. 3-D). *Acauã* compared to *Sf* showed GO terms of DEGs related to ExS, T and U, mainly down-regulated (Fig. 3E), but in a lower number in relation to the other comparisons (Fig. 3D and E) which is in agreement with previous comparisons (Fig. S4-E and S5-E). *Sf* had a greater number of terms enriched for down-regulated genes in relation to *Oeiras* and classified as ExS, FD and HR (Fig. 3F). These conclusions underscore the significance of GO analysis in elucidating the complex interplay of gene expression across diverse coffee genotypes and tissues

For FBs libraries, the most enriched GOs term was HR followed by FD, ExS and EnS with similar patterns independent of genotype but different regarding up-and down-regulated genes (Fig. 3G to I). *Acauã* vs *Oeiras* showed a lower number of GO terms (Fig. 3G) in contrast to the other comparisons (Fig. 3H and I) which was interpreted as a higher transcriptional similarity as previously described (Fig. S4-G and S5-G), and the terms of DEGs were again mainly related to HR followed by ExS (Fig. 3G). On the other hand, the comparisons between *Acauã* or *Oeiras* against *Sf* (Figs. 3H and I, respectively) showed similar patterns, both with a higher number of GO terms up-and down-regulated, mainly enriched for HR followed by FD, ExS, EnS. These results indicated a transcriptional divergence of the floral development for both *Acauã* and *Oeiras* in relation to the *Sf* mutant, which agrees with the phenotypic differences of these coffee genotypes (Fig. 1). Furthermore, these results importantly pointed out that these transcriptional differences are mainly related to GO terms of DEGs related to HR, a term that consistently emerged as the most enriched across all comparisons (Fig. 3)

Similarly, we evaluated the enriched GO terms in the RNA-seq libraries of leaves collected in March or August and for the different coffee genotypes (Fig. 4). In general, the most enriched terms were again related to HR that appeared in all comparisons, independent of tissue and genotype (Fig. 4A to I). In detail, comparing the leaf libraries from March and August for each genotype, *Oeiras* showed GO terms enriched for HR followed by ExS and FD (Fig. 4A), *Acauã* showed terms of DEGs in the same GO classifications but majorly represented by down-regulated genes (Fig. 4B), whereas *Sf* has few DEGs with enriched terms, mostly up-regulated (Fig. 4C). Comparing genotypes in March, *Acauã* vs *Oeiras* and *Acauã* vs *Sf* showed terms of DEGs that are mostly related to ExS and HR (Fig. 4D and 4E, respectively) with a lower number in comparison to *Oeiras* vs *Sf*, which showed a higher number of down-regulated terms enriched for FT (Fig. 4F). In August, the genotype comparisons revealed a congruent trend: *Oeiras* and *Acauã* exhibited greater similarity (evidenced by a lower number of GO terms compared to Sf), HR standing as the most enriched term, followed by FD and ExS (Figs. 4G to I). Notably, the *Sf* genotype displayed fewer enriched GO terms when comparing the two periods (Fig. 4C), which contrasts with its higher GO terms count compared to the other genotypes irrespective of the period (Figs. 4E, F, H, and I). This intriguing observation underscores the *Sf* mutant’s consistent transcriptional pattern in leaves during both March and August, despite the varying environmental conditions of these periods. Such findings further reinforce the hypothesis of transcriptional homeostasis or reduced sensitivity to environmental factors, as supported by data in Table S3 and Fig. S5.

Overall, the conducted GO analysis underscores the interplay of genotype and time in shaping the transcriptional profiles of SAM, FBs, and leaves, particularly evident in the abundance of GO terms associated with HR, as illustrated in Figures 3 and 4. Thus, based on gene annotation (Table S3), we classified the DEGs in the different classes of hormones to know which one was a major factor regulating transcription during floral development or leaf function (Fig. 5 and 6). The analysis was conducted as before, comparing genotypes and tissues, and very interestingly, the most representative hormone was ethylene followed by auxin.

**Fig. 5.**
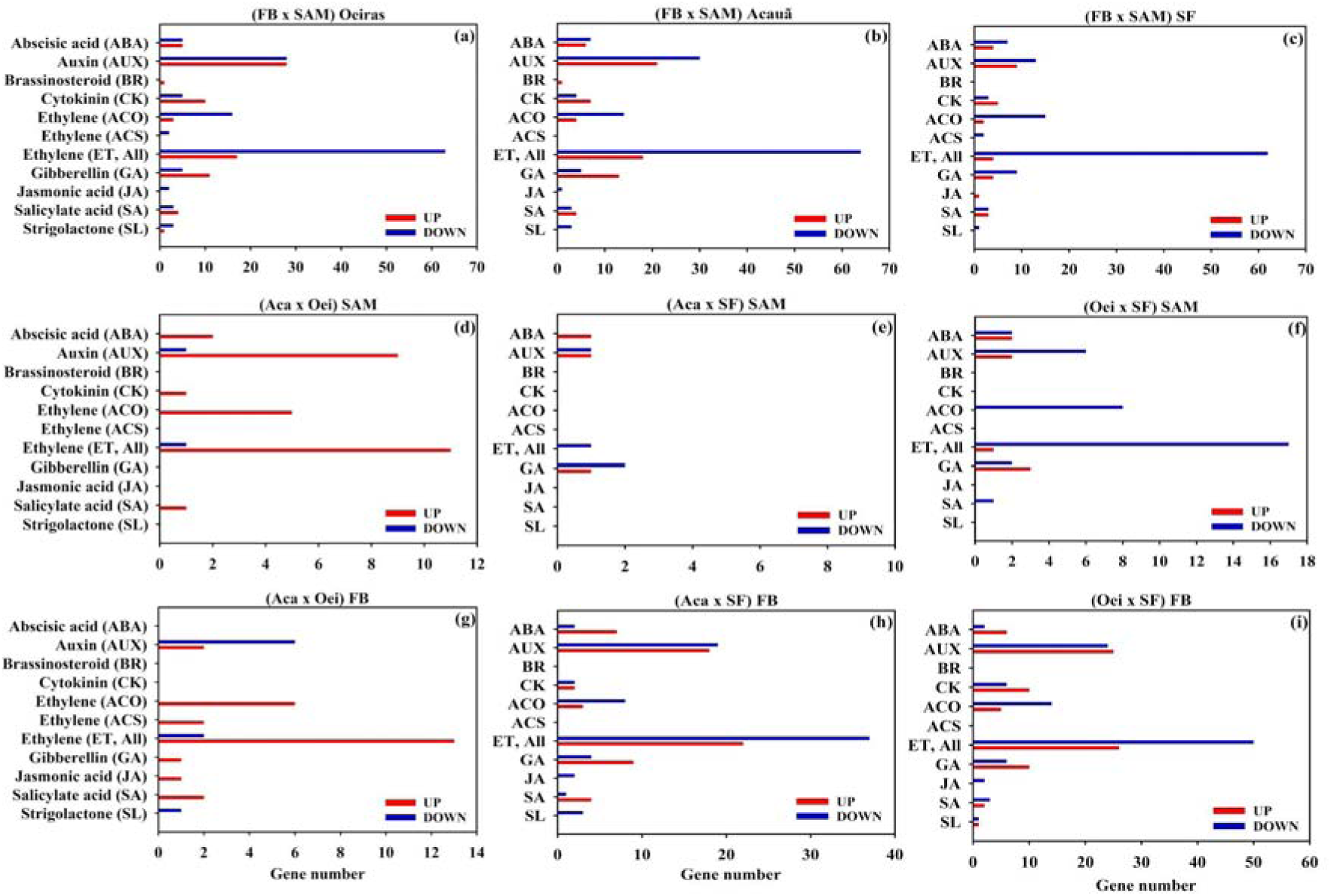
Classification of DEGs in the different classes of hormones found in tissues of the coffee genotypes. The three upper figures show the contrast comparing up-and down regulated genes related to different hormones in different tissues, such as Floral bud (FB) and Shoot Apical Meristem (SAM), of the same genotype: *Oeiras* (Fig. a), *Acauã* (Fig. b) and *Semperflorens* mutant (Fig. c). The contrasts comparing up-and down regulated genes related to different hormones in the same tissue but different genotypes are shown in Figs. d to i. Legends: FB, Floral buds at the G2 stage; SAM, Shoot Apical Meristem; Aca, *Acauã*; Oei, *Oeiras*; SF, *Semperflorens* mutant; UP (in red), number of terms related to UP-regulated genes; DOWN (in blue), number of terms related to DOWN-regulated genes. ABA, Abscisic acid; AUX, Auxin; BR, Brassinosteroid; CK, Cytokinin; ACO, 1-Aminocyclopropane-1-Carboxylic Acid Oxidase which is related to the ethylene pathway; ACS, 1-Aminocyclopropane-1-Carboxylic Acid Synthase which is related to the ethylene pathway; ET (All), all genes found related to the ethylene pathway; GA, Gibberellin; JA, Jasmonic acid; SA, Salicylic acid; SL, Strigolactone. Details of the collected tissues and periods are available in Fig. S1.

**Fig. 6.**
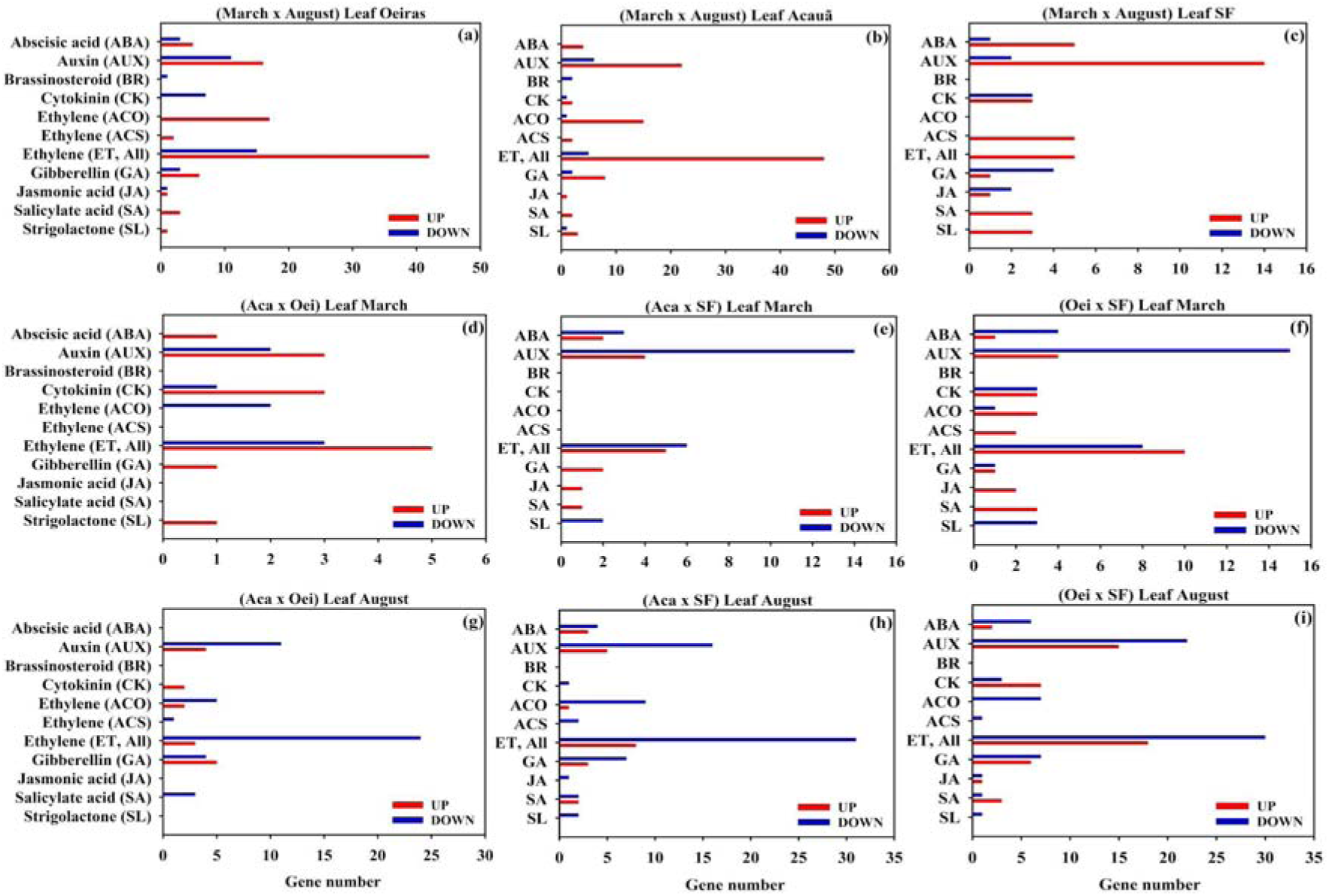
Classification of DEGs in the different classes of hormones found in leaves collected in different periods of coffee genotypes. The three upper figures show the contrast comparing up-and down-regulated genes related to different hormones in leaves collected in March or August of the same genotype: *Oeiras* (Fig. a), *Acauã* (Fig. b) and *Semperflorens* mutant (Fig. c). The contrasts comparing up-and down regulated genes related to different hormones in leaves collected at the same period but different genotypes are shown in Figs. d to i. Legends: Aca, *Acauã*; Oei, *Oeiras*; SF, *Semperflorens* mutant; UP (in red), number of terms related to UP-regulated genes; DOWN (in blue), number of terms related to DOWN-regulated genes. ABA, Abscisic acid; AUX, Auxin; BR, Brassinosteroid; CK, Cytokinin; ACO, 1-Aminocyclopropane-1-Carboxylic Acid Oxidase which is related to the ethylene pathway; ACS, 1-Aminocyclopropane-1-Carboxylic Acid Synthase which is related to the ethylene pathway; ET (All), all genes found related to the ethylene pathway; GA, Gibberellin; JA, Jasmonic acid; SA, Salicylic acid; SL, Strigolactone. Details of the collected tissues and periods are available in Fig. S1.

In detail, the comparisons between SAM and FBs for all genotypes showed a pattern of ethylene and auxin related genes being primarily down-regulated in SAM (Figs. 5A to). Comparing genotypes, ethylene-and auxin-related DEGs are up-regulated in *Oeiras* in comparison to *Acauã* (Figs. 5D), which is tempting to associate with the *Oeiras* early-flowering phenotype (Fig. 1). For SAM libraries, *Acauã* has fewer related DEGs compared to *Sf* and (Figs. 5E) contrasting to *Oeiras* with more down-regulated DEGs (Figs. 5F), which is in accordance with previous results (Figs. 3E and F, Figs. S4-E and -F). For FB library contrasts, *Acauã* vs *Oeiras* showed mostly ethylene-related up-regulated DEGs in *Oeiras* (Fig. 5G) but in a smaller abundance compared to the other comparisons (Fig. 5H and I) indicating transcriptional similarity of FBs between these genotypes. *Acauã* and *Oeiras* compared to *Sf* (respectively, Figs, 5H and I) showed similar patterns with mainly ethylene-and auxin-related DEGs.

For leaf libraries (Fig. 6), the analyses showed a variable pattern of DEGs with *Oeiras* and *Acauã* (respectively, Figs. 6A and B) presenting a higher number of up-regulated DEGs in August related to ethylene, whereas *Sf* showed more DEGs related to auxin and in a smaller number (Fig. 6C). Comparing genotypes, in March a fewer number of DEGs related to ethylene, auxin and cytokinin were found between *Acauã* and *Oeiras* (Fig. 6D) and a higher number was found comparing these genotypes to *Sf* (Figs. 6E and F). In contrast, DEGs are mainly related to auxin and down-regulated in *Sf* compared to both genotypes (Figs. 6E and F). In August, the transcriptional patterns were similar between genotype comparisons with DEGs mainly down-regulated and related to ethylene (Figs. 6G to I).

### 3.6. Differential expression of floral major players could explain coffee phenotypes

To explore genes potentially related to the different coffee flowering phenotypes (Fig. 1), we initially conducted an analysis using a Venn diagram to visualize the shared and unique DEGs resulting from the comparisons between SAM and FB (Figs. A to C in Suppl. doc. Floral genes). We found that 2,700 out of 9,998 DEGs are shared by all genotypes that represent the core of genes distinguishing SAM or FB, 3,408 DEGs shared by *Acauã* and *Oeiras*, whereas 481 and 259 when both genotypes are compared to *Sf*, respectively (Fig. A in Suppl. doc. Floral genes). Genotype-specific DEGs contrasting SAM and FB libraries were also found, including, 1776 for *Oeiras*, 868 for *Acauã* and 506 for Sf (Fig. A in Suppl. doc. Floral genes).

The count of DEGs comparing SAM libraries across genotypes was lower than FB, with 1,805 (Fig. B in Suppl. doc. Floral genes) compared to 7,596 (Fig. C in Suppl. doc. Floral genes). Interestingly, the FB libraries showed the lowest number of DEGs in the comparison of *Acauã* vs *Oeiras* and a much higher abundance when both genotypes were contrasted to *Sf*, with 4,502 DEGs shared between these comparisons (Fig. C in Suppl. doc. Floral genes). This result showed that the transcriptional machinery is highly divergent in the *Sf* mutant compared to other genotypes, aligning with its unusual flowering phenotype (Fig. 1).

Therefore, we seek across the 12,478 DEGs (Suppl. file Barplots and Table S3) for the known players of floral regulatory pathways described in literature and evaluate their expression profiles in the RNA-seq libraries (Suppl. doc. Floral genes). Four DEGs related to hormones were selected: the *ERF027-like* (XM_027245516.1) and *ERF038-like* (XM_027220115.1) related to ethylene responses, which were down-regulated in of *Sf* followed by *Oeiras* and *Acauã* with the highest expression level. In an opposite pattern, an *AUXIN SYNTHETASE* (XM_027261373.1) and a *GIBBERELLIN DIOXYGENASE* (XM_027267466.1) were up-regulated in FBs and SAM of *Sf* compared to other genotypes (Suppl. doc. Floral genes). Regarding DEGs related to floral transition, we found that a gene annotated as *HEADING DATE 3A-like* (*Hd3a-like*; XM_027231814.1), but with identical protein sequence to the floral activator *CaFT1* (Cardon et al., 2022), is much more expressed in FBs of *Sf* compared to the other genotypes and, interestingly, ectopically expressed at SAM of *Sf*. In addition, two homologs of the PEBD family, *CEN-like* (XM_027209634.1) and *MOTHER of FT/TFL1*, together *SQUAMOSA PROMOTER-BINDING-like PROTEIN 3* (XM_027205283.1) and *EARLY FLOWERING 5-like* (XM_027268021.1), presented a similar pattern to *CaFT1* (Suppl. doc. Floral genes).

Floral identity and developmental genes belonging to the MADS-box family of transcription factors and already described in coffee (de Oliveira et al., 2014; Rume et al., 2023) were also explored. For instance, the *CaAP1/CAL* (XM_027240835.1)*, CaAP3* (027235182.1) and *CaPI* (XM_027236968.1) were up-regulated in FBs of the *Sf* mutant compared to other genotypes (Suppl. doc. Floral genes). These patterns were similar to *Hd3a-like* (or *CaFT1*) and in agreement with the positive regulation of ABC-model homeotic genes by the *FT* (Amasino, 2010; Ausin et al., 2005; Teo et al., 2019). On the other hand, homologs of *SVP-like* (XM_027234209.1), *SOC1-like* (XM_027229901.1) and *AGL27* (XM_027241945.1), showed an opposite trend being down-regulated in FBs of *Sf* compared to other genotypes, in agreement with previous works (de Oliveira et al., 2014; Rume et al., 2023).

Other important genes related to floral development were also evaluated: *ACIDIC ENDOCHITINASE-like* (XM_027239564.1) and *VIN3-like PROTEIN 2* (XM_027235309.1) that were more expressed in FBs of the mutant *SF*, whereas *LIGHT-DEPENDENT SHORT HYPOCOTYLS 10-like* (XM_027209174.1), *NRT1/ PTR FAMILY 2.11-like* (XM_027266104.1) and *COLD AND DROUGHT-REGULATED PROTEIN CORA*-*like* (XM_027263410.1) were less expressed in FBs of the mutant *SF* compared to other genoptypes (Suppl. doc. Floral genes).

## 4. DISCUSSION

### 4.1. Floral development and flowering time plasticity in *Coffea*

An interesting aspect of coffee flowering is that *C. arabica* has a poor genomic variability due a unique event of polyploidization that originated the species (Lashermes et al., 1999; Scalabrin et al., 2020) between 10,000 and 665,000 years ago (Cenci et al., 2012; Q. Yu et al., 2011; Salojärvi et al., 2023). However, somehow coffee genotypes present plasticity with different characteristics related to floral development that are useful for breeding programs. For example, *Acauã* genotype is a late-flowering, *Oeiras* early-and *Semperflorens* present a continuous flowering (Antunes, 1960; Carvalho & Krug, 1952) Carvalho et al., 2008). This plasticity appears to be related to the regulation of the reproductive or circadian cycle, as demonstrated by various studies in different crops (Komaki & Sugimoto, 2012; Paajanen et al., 2021; Xu et al., 2023). Nevertheless, the floral development patterns of contrasting genotypes and their associated regulatory pathways, especially regarding flowering time, have not been reported in detail yet in *C. arabica* (López et al., 2021).

Another unexplored aspect in this work is the relationship between the reproductive development of coffee plants and endogenous/environmental stimuli. The coffee cycle is complex, presenting an unusual biennual cycle for crops (López et al., 2021). For example, in Brazil, the coffee cycle is described as highly influenced by cold temperatures and water availability to activate and synchronize flowering, which significantly impacts productivity and quality (DaMatta et al., 2007; DaMatta & Ramalho, 2006). However, despite the floral transition of coffee being related to conserved pathways, such as the florigen CaFT1 and small RNAs (Cardon et al., 2022; Ribeiro et al., 2024), the molecular link between coffee flowering and environmental cues has not been established yet (Cardon et al., 2022; López et al., 2021).

Thus, we confirmed the contrasting phenotypes, describing in detail their reproductive cycle from the beginning of floral bud differentiation until the flowering, including the number of observed anthesis events of each genotype (Fig. 1). The coffee phenotypic plasticity was associated with the differential sugar status during development (Fig. 2) and the differential expression of hundreds of genes, mainly related to environmental response, floral development and hormonal regulation (Figs. 3 to 6 and Table S3). Studies of reproductive development directly impact yield and quality (Xu et al., 2023) and are crucial for ensuring food production in face of climate changes, especially in crops with low genetic variability. These aspects along with our results are discussed below.

### 4.2. Genes involved in sugar metabolism and floral development

Sugars are the main product of photosynthesis and are mobilized to different plant organs for growth tissue differentiation. Previous research indicates that sucrose function primarily in the leaf phloem to promote the transcription of the florigen *FT*, whereas Trehalose-6-phosphate acts in the SAM to promote the signaling pathway downstream of the flowering process (Cho et al., 2018). The role of sugar in the flowering time has been extensively studied since sugar represents energy and building material for the metabolic processes.

Sucrose promotes flowering in various species (Bernier et al., 1993). Transcriptome analysis using mature leaves in maize (*Zea mays* L. spp *mays*) sampled at the floral transition stage has shown that expression levels of several key genes involved in starch and sucrose metabolism are altered in the *id1* mutant (Coneva et al., 2012). These findings suggested that the balance between transitory starch and sucrose is crucial for controlling flowering time. Increased level of endogenous sucrose also enhances flowering in tomatoes (*Solanum Lycopersicum* L.) (Micallef et al., 1995).

Flowering events were more frequent in the mutant *Sf* compared to *Acauã* and *Oeiras*, with *Oeiras* displaying a higher number of flowering events than *Acauã* (Fig. 1). Notably, in *Acauã*, sucrose content in leaves appears crucial in proximity to the primary flowering event in September, which is evident from the escalating levels observed from June through August (Fig. 2). In contrast, for *Sf* and *Oeiras*, sucrose content is distributed more evenly across multiple periods due to their more frequent flowering events. While total sugars experienced an upswing in May for *Oeiras*, this trend was not mirrored in *Acauã* and *Sf*. Instead, there was a discernible peak in July, followed by a decrease prior to anthesis in August (Fig. 2). The pattern for reducing sugars exhibited an upward trajectory, yet no discernible distinctions emerged among the coffee genotypes (Fig. 2).

The RNA-seq analysis unveiled a substantial number of both up and down-regulated genes intricately linked to sugar metabolism (Figs. 3, 4, Table S3, Suppl. doc. Floral genes). Notably, these encompassed genes associated with enzymatic processes involved in sugar synthesis and cleavage, as well as those governing intracellular sugar transport (Figs. 3, 4 and Table S3).

The flowering events were higher in *Sf* than *Acauã* and *Oeiras*, whereas *Oeiras* presented more flowering events than *Acauã* (Fig. 1). It seems that sucrose content in *Acauã* is needed close to the main flowering event in September and for that reason, we observed an increase from June to August, whereas for *Sf* and *Oeiras*, sucrose content is not concentrated in a specific period because flowering events are more frequent. Total sugars increased in May for *Oeiras* but not for *Acauã* and *Semperflorens*, showing a peak in July, decreasing before anthesis in August (Fig. 2). Reducing sugars showed an increased pattern but no differences were observed between the coffee genotypes (Fig. 2).

In the Gene Ontology analysis of leaves, FB and SAM transcriptomes, we found some interesting contrasts showing that sugar is involved in those tissues, possibly as signaling of flowering time. A higher number of DEGs were found in leaves than FB and SAM because the photosynthetic process is superior in leaves (Table S3 and Suppl. doc. Floral genes). The expression of two genes as candidates for starch biosynthesis in *Sf* in March (glucose-6-phosphate/phosphate translocator chloroplastic-like and glucose-1-phosphate adenylyltransferase large subunit 1-like) (Table S3 and Suppl. doc. Floral genes). Two candidate genes as sucrose transporter in August (*SUGAR TRANSPORT PROTEIN 13-like* and *HEXOSE CARRIER PROTEIN* HEX6-like) (Table S3 and Suppl. doc. Floral genes). The expression of those genes is congruent with the high starch content observed in *Sf* in March and the increased level of sucrose for *Acauã* in August (Fig. 2), respectively.

During the analysis of leaves, FB, and SAM transcriptomes through Gene Ontology (GO), intriguing distinctions emerged, hinting at sugar’s potential role in these tissues, possibly as a signaling mechanism for regulating flowering time. Notably, a more pronounced number of DEGs surfaced in leaves as compared to FB and SAM (Table S3 and Suppl. doc. Floral genes). This disparity can be attributed to the heightened photosynthetic activity occurring within leaves. The expression profiles of two specific genes are showcased as prospective candidates responsible for starch biosynthesis in *Sf* during March (Table S3 and Suppl. doc. Floral genes). These genes are the *GLUCOSE-6-PHOSPHATE/PHOSPHATE TRANSLOCATOR CHLOROPLASTIC-like* and *GLUCOSE-1-PHOSPHATE ADENYLYLTRANSFERASE LARGE SUBUNIT 1-like* (Table S3 and Suppl. doc. Floral genes). Remarkably, the expression patterns of these genes align with the observed high starch content in *Sf* during March, as well as the elevated sucrose levels seen in *Acauã* during August (Fig. 2 and Suppl. doc. Floral genes). This congruence between gene expression and biochemical observations highlights the potential functional relevance of these genes in regulating sugar-related processes within the coffee genotypes.

### 4.3. Hormonal regulation in coffee flowering

Despite the absence of any effective models for the hormonal controls of flowering, numerous studies have demonstrated the individual effect of plant hormones in flowering (Izawa, 2021). Changes in ethylene levels can either delay or promote flowering, as observed in Rice (*Oryza sativa* L.) (Wang et al., 2013), *Arabidopsis* (*Arabidopsis thaliana*) (Achard et al., 2007), Pineapple (A*nanas comusus* L.) (Trusov & Botella, 2006), and Roses (Meng et al., 2014). In coffee, recent research suggests that ethylene can play an important role in flowering, because rehydrating droughted plants can increase ethylene levels and ethylene sensitivity, regulating coffee anthesis (Lima et al., 2021).

For this study we evaluated those expressed genes related to ethylene production, with the majority of these genes involved in biosynthesis pathway (*AMINOCYCLOPROPANE-1-CARBOXYLATE-like*), and receptors for signaling cascade (*ETHYLENE-RESPONSE TRANSCRIPTION FACTOR RAP2-12*-*like*, *ETHYLENE-RESPONSIVE TRANSCRIPTION FACTOR 1B-like*, *EIN3-BINDING F-BOX PROTEIN 1-like*, *ETHYLENE-RESPONSIVE TRANSCRIPTION FACTOR 4-like*, and *ETHYLENE-RESPONSIVE TRANSCRIPTION FACTOR 5-like*) (Table S3 and Suppl. doc. Floral genes). In the used genome annotation, Cara_1.0, there are sixty-three *1-AMINOCYCLOPROPANE-1-CARBOXYLATE OXIDASE (ACO)* and eleven *1-AMINOCYCLOPROPANE-1-CARBOXYLATE SYNTHASES (ACS)*. Twenty-eight ACO genes were found to be differentially expressed in at least one contrast whereas two ACS were DE in at least one contrast (Tables S3 and Suppl. doc. Floral genes). The ACS and ACO enzymes, which control the rate-limiting steps of ethylene biosynthesis, are encoded by multigene families in other species (Arraes et al., 2015; Seymour et al., 2013; Yamagami et al., 2003; Yang & Hoffman, 1984). Possibly the difference in the chemical composition of cells, tissues and organs can influence the specificity of the expression of homologous genes and, in coffee, the expression of *ACS* and *ACO* genes varies according to the organ and the environmental condition (Lima et al., 2021; Ságio et al., 2014; Santos et al., 2022). A regulatory mechanism for the ethylene biosynthesis pathway associated with anthesis has been suggested, showing a repression of the three *ACO* genes in the roots during the dry season. However, in floral buds, these gene expressions are independent of soil water content, suggesting greater control of ethylene production in the reproductive organs during the final stages of flowering (Santos et al., 2022).

Connected to this model of transcriptional regulation of ethylene biosynthesis genes, it has been demonstrated that ethylene levels in coffee roots are reduced during the dry season and increased after rehydration, preceding anthesis (López et al., 2022). In this study, the ethylene level was measured in leaves and floral buds of the coffee genotypes *Oeiras*, *Acauã* and *Sf* (Fig. 2E and 2F). In general, leaves ethylene levels in the *Acauã* and *Oeiras* were increased in the wet season and reduced in the dry season, while the opposite occurred in floral buds. These variations indicate the existence of different ethylene levels between buds at different stages of development, as highlighted in our study, which is in agreement with the findings of Schuch et al., (1992). Furthermore, it has been suggested that the significance of ethylene for coffee flowering is related to its ability, after breaking the latent state of G4 floral buds, to promote uniform anthesis (Lima et al., 2021; López et al., 2022). However, we believe that ethylene is involved not only in the anthesis process but in all the coffee flowering processes. This is evidenced in our RNA-seq analysis, which displayed a high number of UP and DOWN-regulated genes in the biosynthesis and signaling pathway of ethylene from undifferentiated tissue in SAM to floral buds in the beginning of development (Fig. 5 and Suppl. doc. Floral genes). Our hypothesis could explain the frequent flowering observed in the mutant coffee genotype *Sf*, which does not seem to depend on exogenous factors such as drought and rain. In this case, it is possible that flowering in *Sf* is more related to an endogenous regulation with active participation of the hormonal pathway governed mainly by ethylene.

In the DEGs analysis, the most remarkable for all evaluated contrasts in leaves, FB, and SAM, is the amount of UP and DOWN-regulated genes associated with hormonal regulation (Figs. 5 and 6). While ethylene is implicated in coffee flowering time regulation, other hormones such as gibberellins, auxins, strigolactones, cytokinins, and abscisic acid may also play roles as signaling in response to the environmental cues or endogenous stimulus. Previous studies have shown that Abscisic Acid (ABA) increases during the dry period and is associated with coffee floral development in “latent state”, with plant rehydration decreasing ABA and increasing ethylene and Gibberellin (GA) levels (Browning, 1973; López et al., 2022). In *Arabidopsis*, GA in leaf and shoot apex tissues can promote flowering under long-day conditions, whereas apex-specific depletion of GA results in non-flowering phenotypes under short-day conditions (Porri et al., 2012). On the other hand, a genetic link between GA and Jasmonic Acid (JA) was established, because DELLA protein can enhance the activity of TARGET OF EAT1 (TOE1) and TOE2 indirectly via sequestration of JAZ (jasmonate-ZIM domain). As a result, DELLA degradation by GAs frees multiple repression sites in the *FT* promoter, controlling flowering time in *A. thaliana* (Izawa, 2021). Exogenous cytokinin can promote flowering under short-day conditions. *TSF* (another florigen in *A. thaliana*), *FD*, and *SOC1* are required for cytokinin-mediated flowering (D’Aloia et al., 2011).

UP and DOWN-Regulated genes associated with auxin reveal that this hormone can also play a significant role from floral initiation to floral buds development in coffee trees (Fig 5). Under natural conditions, auxin inhibits flowering in general (Achard et al., 2007; Kęsy et al., 2010). Nonetheless, it has been reported that auxin application can inhibit flowering by blocking florigen translocation, or can induce the floral buds development after the florigen reaches the shoot apical meristem (Salisbury, 1955). Thus, auxins play a role in specifying the number and identity of floral organs (Cheng & Zhao, 2007). Furthermore, auxin application in pineapple promoted an increase in endogenous ethylene levels and induced flowering (Burg & Burg, 1966). In our results, the central role that ethylene plays on coffee flowering seems to be supported, primarily, by auxin in different floral development phases and, secondarily, by ABA and GA in later stages, mainly related to anthesis.

### 4.4. Candidate genes involved in coffee flowering time in response to endogenous and external stimulus

The timing of flowering is determined by both endogenous genetic components and various environmental factors, such as day length, temperature, and stress. Plants respond to these stimuli by a signaling cascade, directing to the transition of the vegetative to reproductive phase. In this context, numerous interconnected genes form regulatory networks for specific functions. Our study identified thousands of genes potentially associated with different functions in the flowering timing process. These genes help explain the differences in flowering time among the contrasting coffee genotypes. We selected a subset of 250 DEGs UP and DOWN-Regulated genes related to floral identity and flower development as candidates to elucidate these differences (Table S3 and Suppl. doc. Floral genes).

In the classification of Gene Ontology (GO) for flower development, we can note some differences between contrasting coffee genotypes (Fig. 3 and 4). Within the context of GO classification pertaining to flower development, discernible distinctions emerge when comparing contrasting genotypes. In the case of FB, there are several more up-regulated genes for *Oeiras* compared to *Acauã* in March, even though they were phenotypically similar in terms of flower development at this point (Fig. 1). The higher number of DEGs in *Oeiras* than *Acauã* could suggest a transcriptional difference related to their flowering pattern (Table S3 and Suppl. doc. Floral genes). The pattern of up and down-regulated genes for flower development comparing *Acauã* and *Oeiras* against *Sf* is similar. This difference seems particularly pronounced in the SAM because only up-regulated genes of flower development were observed when comparing the SAM of *Oeiras* and *Acauã* (Fig. 3, Table S3 and Suppl. doc. Floral genes).

As expected, there are more terms enriched in FB than in SAM contrasts (Fig. 3). In the pursuit of unraveling the genes responsible for orchestrating flower development across leaves, FB, and SAM, we focused on some known players exhibiting the highest expression levels (Tables S3, GO and barplot figures, Suppl. doc. Floral genes). For example, the protein HEADING 3A-like is a homolog of FT in rice, known to promote flowering in short days. Due to the high level of expression observed in *Sf* FB compared with *Acauã* and *Oeiras*, we can speculate that this up-regulation could explain why *Sf* are different regarding its flowering time (Tamaki et al., 2007; Taoka et al., 2011) (Suppl. doc. Floral genes). Similarly results were found for other genes such as the transcription factor CAULIFLOWER (CAL) described as a probable transcription factor that promotes early floral meristem identity in synergy with APETALA1 (AP1), FRUITFULL (FUL), and LEAFY (LFY) (Tables S3, GO and barplot figures, Suppl. doc. Floral genes). CAL is required subsequently for the transition of an inflorescence meristem into a floral meristem and it seems to be partially redundant to the function of APETALA1 (X. F. Li et al., 2000).

The *CAL* primarily specifies the floral meristem identity in a redundant manner to *AP1* and *FUL* genes that have also functions in several processes, such as the development of carpel structures and specification of inflorescence meristem (IM) identity (Ferrándiz et al., 2000). *CAL* is highly expressed in FB of *Sf*, possibly regulating early flowering in this genotype (Tables S3, GO and barplot figures, Suppl. doc. Floral genes). The MADS-box protein *SOC1-like* is more highly expressed in FB of *Oeiras* than in *Acauã* and *Sf* (Tables S3, GO and barplot figures, Suppl. doc. Floral genes), potentially explain the difference in flowering time of *Oeiras* and *Acauã*, because this is a transcription activator acting in flowering time control. When associated with *AGL24, CAL* mediates the effect of gibberellins on flowering under short-day conditions, and regulates the expression of *LEAFY (LFY)*, which links floral induction and floral development (Liu et al., 2008). The K-domain of a blueberry-derived *SOC1-like* gene promotes flowering in tobacco (Song et al., 2013). Overexpression of *ZmSOC1* resulted in early flowering in *Arabidopsis* through increasing the expression of *AtLFY* and *AtAP1*. Overall, these results suggest that *ZmSOC1* is a flowering promoter in *Arabidopsis* (Zhao et al., 2014).

The gene *VIN3-like protein 2* is highly expressed in FB of *Sf* and represents other candidate gene involved in coffee floral regulation in this genotype (Tables S3, GO and barplot figures, Suppl. doc. Floral genes). This gene may be involved in both the vernalization and photoperiod pathways by regulating gene expression transcription and promoting flowering in non-inductive photoperiods (Sung et al., 2007).

The *MOTHER of FT* protein and *TFL1 homolog 1-like* are among the most expressed in *Sf* compared to *Acauã* and *Oeiras* (Tables S3 and Suppl. doc. Floral genes), being a candidate for regulating flowering patterns in this genotype. These genes regulate identities of the determinate and indeterminate meristems, and ultimately affect flowering time and plant architecture (Yu et al., 2019). MOTHER of FT and TFL1 functions as floral inducers and may act redundantly in the determination of flowering time in *Arabidopsis* (Yoo et al., 2004). Additionally, six candidates’ genes were identified acting in the autonomous pathway in the flowering time control (Tables S3, GO and barplot figures, Suppl. doc. Floral genes). Finally, the *EARLY FLOWERING 5-like* protein was selected, a gene that has been described as a flowering repressor in *Arabidopsis* (Noh et al., 2004). However, it is possible that due to its expression pattern in all the evaluated tissues of coffee genotypes (Leaves, FB and SAM), this gene could be acting as a flowering promoter, especially in *Sf* (Tables S3, GO and barplot figures, Suppl. doc. Floral genes). At this point, we cannot explain how this could happen, it provides an intriguing proposal for future studies.

## 5. CONCLUDING REMARKS

Differences in the coffee floral development and flowering-time of contrasting among coffee genotypes appear to be associated with sugar metabolism and genes mainly related to hormonal regulation, floral development and environmental stimuli. Specifically, the *Sf* mutant showed higher levels of starch and differential expression of key factors within regulatory floral pathways compared to the other genotypes. Moreover, the analysis of differential expression in leaves collected across in different periods showed fewer transcriptional differences for *Sf* than for *Oeiras* and *Acauã*, suggesting that this mutant genotype is less influenced by environmental cues. This apparent lack of transcriptional sensitivity to the environment may be associated with its continuous flowering phenotype.

Examining the gene expression dynamics associated with key hormones like ethylene, auxin, and gibberellin, alongside those linked to floral pathways, reveals a compelling correlation with the delayed and early-flowering traits exhibited by *Acauã* and *Oeiras*, respectively. The expression patterns of genes related to hormones, mainly ethylene, auxin and gibberellin, and floral pathways, correlate with the late-and early-flowering phenotypes of *Acauã* and *Oeiras*, respectively. Very interestingly, these genes also appear deregulated in the *SF* mutant, for example, the florigen *CaFT1* is ectopically expressed in SAM and FBs which could explain the continuous meristem production and floral transition. Thus, in summary, our study demonstrates contrasting floral development between coffee genotypes (Figs. 1 and 2), primarily regulated by differences in floral-ethylene-and auxin-related pathways (Figs. 3 to 6).

A recent work has explored the structure of *C. arabica* chromosomes revealing aberrations and exchanges related to genetic diversity (Scalabrin et al., 2024). However, future studies should explore the causes of differential expression between genotypes and identify the main pathways resulting in phenotype diversity. In this way, it would be enlightening to explore the role of transposons and epigenetic contribution for transcriptional regulation in coffee. Moreover, the functional characterization of unknown genes, which occur in a higher number in the coffee genome, and a deeper exploration of *SF* development, could reveal new genes or pathways controlling floral development. In conclusion, our work contributes to understanding the divergence of floral development in crops with low genetic variability directing breeding programs and future research towards the control of floral development and production improvement.

## AUTHORS’ CONTRIBUTIONS

M.E.L., R.R.O., D.Z. and A.C-J. conceptualized the project; M.E.L. and L.M.A. conducted all experiments and data research; T.H.C.R. and D.Z. supported for bioinformatics analyses; M.E.L., R.R.O., L.M.A. and I.S.S. discussed all data and analysis; R.R.O. and A.C-J supervised experiments and analyses; M.E.L., R.R.O., L.M.A. and I.S.S. wrote the manuscript; all co-authors corrected and contributed to writing of the final manuscript version.

## ACKNOWLEDGEMENTS

The authors thank the Federal University of Lavras (UFLA/Brazil), the members of the Laboratory of Plant Molecular Physiology (LFMP, UFLA/Brazil) and Agricultural Research Center (USDA/ARS) for structural support of experiments and analysis, and the funding agencies for all the support provided

## STATEMENTS AND DECLARATIONS

The authors declare that there are no conflicts of interest.

## DATA AVAILABILITY

Main data supporting the findings of this study are available within the paper and within its supplementary materials published online. Raw data for the RNA-seq runs can be obtained through BioProject ID PRJNA1091177. The respective SRA accessions are available in Table S1.

